# Ant collective behavior is heritable and shaped by selection

**DOI:** 10.1101/567503

**Authors:** Justin T. Walsh, Simon Garnier, Timothy A. Linksvayer

## Abstract

Collective behaviors are widespread in nature and usually assumed to be strongly shaped by natural selection. However, the degree to which variation in collective behavior is heritable and has fitness consequences -- the two prerequisites for evolution by natural selection -- is largely unknown. We used a new pharaoh ant (*Monomorium pharaonis*) mapping population to estimate the heritability, genetic correlations, and fitness consequences of three collective behaviors (foraging, aggression, and exploration) as well as body size, sex ratio, and caste ratio. Heritability estimates for the collective behaviors were moderate, ranging from 0.17 to 0.32, but lower than our estimates for the heritability of caste ratio, sex ratio, and the body size of new workers, queens, and males. Moreover, variation among colonies in collective behaviors was phenotypically correlated, suggesting that selection may shape multiple colony collective behaviors simultaneously. Finally, we found evidence for directional selection that was similar in strength to estimates of selection in natural populations. Altogether, our study begins to elucidate the genetic architecture of collective behavior and is one of the first studies to demonstrate that it is shaped by selection.

## Introduction

Collective behavior, defined as behaviors of groups of individuals that operate without central control (Gordon 2014, 2016), is ubiquitous in nature. Examples include predator avoidance in schools of fish, the migration of flocks of birds, and nest building in social insects. Increasingly, researchers have documented patterns of variation in collective behavior between groups (i.e. describing collective or group personality; Gordon 1991; Gordon et al. 2011; Jandt et al. 2014; Bengston & Jandt 2014; Wright et al. 2019) with a goal of understanding the evolutionary causes and consequences of variation in collective behavior. However, the degree to which collective behaviors are heritable and how genetic variation contributes to population-level variation in individual and collective behaviors remain largely unknown. Furthermore, it is often assumed that collective behavior and other group-level traits, like individual behavior and other individual-level traits, are strongly shaped by natural selection. However, little is actually known about the fitness consequences of variation in collective behaviors, or group-level traits more generally (Gordon 2013, 2016; Wright et al. 2019).

Given that trait variation must be heritable in order for the trait to respond to selection and evolve over time, quantifying heritability is a crucial first step in studying trait evolution (Falconer & Mackay 1996; Lynch & Walsh 1998). Previous studies in ants, honey bees, and sticklebacks suggest that collective behaviors and other group-level traits are heritable (Linksvayer 2006; Hunt et al. 2007; Wark et al. 2011; Gordon 2013; Greenwood et al. 2015; Friedman & Gordon 2016). Additionally, candidate gene studies have linked allelic variation to variation in collective behavior, providing further evidence that collective behavior is heritable (Krieger 2005; Wang et al. 2008; Wang et al. 2013; Tang et al. 2018). Although numerous studies have examined the genetic architecture of group-level traits in honey bees (Rinderer et al. 1983; Collins et al. 1984; Milne 1985; Moritz et al. 1987; Bienefeld & Pirchner 1990; Pirchner & Bienefeld 1991; Harris & Harbo 1999; Boecking et al. 2000; Hunt et al. 2007), we know little about the genetic architecture or the evolution of collective behavior and other group-level traits in other group-living species.

Another key factor affecting the relationship between genotype, phenotype, and evolutionary response to selection is the pattern of genetic correlations, i.e. the proportion of variance that two traits share due to genetic causes. Genetic correlations can either accelerate or slow down the rate of evolutionary response to selection, depending on the direction of the correlation relative to the direction of selection on the traits (Lynch & Walsh 1998; Wilson et al. 2010). Understanding genetic correlations is especially important for the study of behavioral evolution since behaviors are often thought to be correlated with each other, forming sets of tightly linked traits that are often described as behavioral syndromes (Sih et al. 2004; Dochtermann & Dingemanse 2013). Although genetic correlations have been estimated for individual-level behaviors (reviewed by van Oers et al. 2005), few studies have examined genetic correlations between collective behaviors or other group-level traits in social insects (except for honey bees; Milne 1985; Bienefeld & Pirchner 1990; Boecking et al. 2000).

The genetic architecture of group-level traits such as collective behavior is likely more complex than the genetic architecture of individual-level traits, because variation in group-level traits arises from phenotypic and genotypic variation within and among groups (Linksvayer 2006, 2015; Bijma et al. 2007a, 2007b; McGlothlin et al. 2010; Gempe et al. 2012). For example, the genotype of each individual may influence its activity rate, which in turn may affect interactions among group members and the collective performance of the group. Thus, group-level traits depend on the genotypes of multiple interacting individuals, just as individual-level traits that are affected by social interactions, as considered in the interacting phenotypes framework (Moore et al. 1997; McGlothlin et al. 2010). Indeed, previous honey bee studies quantifying heritability for colony-level performance traits such as honey yield have treated colony performance as a worker trait that is influenced by the expected genotype of workers and also potentially influenced by the genotype of the queen (i.e. through a maternal genetic effect) (Bienefeld and Pirchner 1990; Bienefeld and Pirchner 1991; Bienefeld et al. 2007; Brascamp et al. 2016).

The rate and direction of a trait’s potential evolutionary response to selection also depends on the pattern of natural selection acting on the trait. Knowledge of the fitness consequences of trait variation allows researchers to characterize the type (e.g., directional, stabilizing, or disruptive) and strength of natural selection acting on a trait (Lande & Arnold 1983; Arnold & Wade 1984; Janzen & Stern 1998; Morrissey & Sakrejda 2013). Many studies have estimated the fitness consequences of individual-level behavioral variation (reviewed by Smith & Blumstein 2008), but the consequences of group-level variation have received relatively little attention (but see Wray et al. 2011; Modlmeier et al. 2012; Gordon 2013; Blight et al. 2016a; Blight et al. 2016b).

Social insects are well-established models for studying collective behavior. Well-studied collective behaviors include nest choice in acorn ants (*Temnothorax* spp.; Möglich 1978; Franks et al. 2003; Pratt 2017), nest defense and hygienic behavior in honey bees (*Apis mellifera*; Spivak 1996; Breed et al. 2004; Evans & Spivak 2010), and the regulation of foraging in pharaoh ants (*Monomorium pharaonis*; e.g. Beekman et al. 2001; Sumpter & Beekman 2003; Robinson et al. 2005) and harvester ants (*Pogonomyrmex barbatus*; e.g. Gordon 2002; Greene & Gordon 2007; Gordon et al. 2007; Gordon et al. 2011; Gordon 2013). The collective behavior of colony members also shapes colony productivity and the relative investment in workers versus reproductives (i.e. caste ratio) and reproductive males versus queens (i.e. sex ratio). Social insect sex ratio and caste ratio have long served as important models for empirically testing predictions from inclusive fitness theory regarding predicted conflicts between queens and workers over sex ratio and caste ratio (Trivers & Hare 1976; Reuter & Keller 2001; Mehdiabadi et al. 2003; Linksvayer 2008; Bourke 2015;). However, despite this long-term intense interest in the evolution of colony-level traits, empirical evidence is scarce about the key parameters governing the evolution of these traits, especially for ants. Indeed, while recent molecular studies have begun to characterize the genomic, transcriptomic, and epigenetic differences between species, between castes within a species, and between individual workers (Friedman & Gordon 2016; Gospocic et al. 2017; Warner et al. 2017; Chandra et al. 2018; Walsh et al. 2018), little is known about the genetic architecture of collective behavior, caste ratio, and sex ratio (Linksvayer 2006). Similarly, while it is clear that colony-level phenotypes can be shaped by patterns of selection within- and between-colonies (Owen 1986; Moritz 1989; Ratnieks & Reeve 1992; Tsuji 1994, 1995; Banschbach & Herbers 1996; Tarpy et al. 2004; Gordon 2013), few studies have attempted to empirically quantify patterns of selection acting on social insect traits.

In this study we used a genetically and phenotypically variable laboratory population of pharaoh ants (*Monomorium pharaonis*). Such a mapping population has proven powerful to elucidate the genetic architecture of a range of traits, including behavioral traits, in mice, rats, and fruit flies (Hansen & Spuhler 1984; Mott et al. 2000; Valdar et al. 2006; King et al. 2012). We first assayed colony-level foraging, aggression, and three measures of exploration using three replicate sub-colonies of 81 distinct colony genotypes of known pedigree (243 replicate sub-colonies total). Collective behaviors are defined as emergent behaviors of groups of individuals that operate without central control, through local interactions (Gordon 2014, 2016). We consider the behaviors of foraging, exploration, and aggression to be collective because all three consist of emergent patterns of workers operating at least in part through local interactions, either through direct antennal contact with other workers or through the influence of pheromones (Adler & Gordon 1992; Gordon & Mehdiabadi 1999; Gordon 2002, 2010; Greene & Gordon 2007; Pinter-Wollman et al. 2013; Kleineidam et al. 2017). In many social insects, foragers are stimulated to begin foraging through interactions with other foragers/scouts returning to the nest (e.g. Gordon 2002; Fernandez et al. 2003; Pinter-Wollman et al. 2013). Both foraging and exploratory behavior are often regulated through the use of trail pheromones (e.g. Fourcassie & Deneubourg 1994; Jackson & Châline 2007). During aggressive responses to threats, workers are often recruited via the use of alarm pheromones (Loftqvist 1976; Blum 1996) or through social interactions with other workers (Kleineidam et al. 2017). We also chose these collective behaviors because they are linked to colony success in other social insects, including other species of ants (Wray et al. 2011; Modlmeier et al. 2012; Blight et al. 2016a; Blight et al. 2016b). Furthermore, we measured colony productivity, caste and sex ratio, and worker, gyne, and male body size. We used the known pedigree of colonies in our mapping population, together with trait measurements in an animal model framework, to estimate the heritability of and genetic correlations between all traits. Finally, we estimated the strength and pattern of selection acting on all the measured phenotypes in the laboratory.

## Materials and Methods

### (a) Background and overall design

All *M*. *pharaonis* colonies used in this study were reared in the lab and derived from eight initial lab stocks, collected from eight different locations across Africa, Asia, Europe, and North America (Schmidt 2010; Schmidt et al. 2010). Specifically, the eight initial stocks were systematically intercrossed for nine generations in order to create a mapping population that was initially designed to be analogous to the mouse heterogeneous stock (Mott et al. 2000; Valdar et al. 2006). After nine generations of intercrossing, each colony in the resulting mapping population is expected to contain a unique mixture of alleles from the eight initial stocks (Pontieri et al. 2017) (**Supplemental figure 1**). We maintained all colonies at 27 ± 1 °C and 50% relative humidity on a 12:12 hour light:dark cycle. We split each colony (henceforth “colony genotype”) into three equally-sized replicates (henceforth “colony replicate”) by emptying the colony genotypes into plastic bowls, gently mixing the queens, workers, and brood, and using tea spoons to scoop them into three new colony containers. Next, we manually counted all individuals within the colony replicates and adjusted the numbers accordingly so that all colony replicates initially consisted of 4 queens, 400 ± 40 workers, 60 ± 6 eggs, 50 ± 5 first instar larvae, 20 ± 2 second instar larvae, 70 ± 7 third instar larvae, 20 ± 2 prepupae, and 60 ± 6 worker pupae. These numbers represent a typical distribution of developmental stages in a relatively small *M. pharaonis* colony (Warner et al. 2018). Except when starving the colony replicates (see below), we fed all colony replicates twice per week with an agar-based synthetic diet (Dussutour & Simpson 2008) and dried mealworms. The colony replicates always had access to water via water tubes plugged with cotton and nested between two glass slides (5 cm x 10 cm). We kept all colony replicates in a plastic colony container (18.5 cm x 10.5 cm x 10.5 cm) lined with fluon and surrounded by a moat of oil to prevent the workers from escaping the box.

After setting up the colony replicates, we gave them two weeks to acclimate to the new conditions before conducting behavioral assays. We fed the colony replicates twice per week except for the week prior to the exploratory and foraging assays during which we starved the colony replicates so that they would be motivated to explore and forage. We conducted the exploratory and foraging assays during the third week and the aggression assays during the fourth week after setting up the replicate colonies.

### (b) Behavioral observations

#### (i) Exploratory assay

We conducted the exploratory assay after the colony replicates had been starved for six days. We assayed the exploratory behavior of both entire colony replicates and groups of five foragers. We conducted the assay inside a filming box with white LED lights arranged along the walls and a camera mounted on the top to film the arena from above (**Supplemental figure 2A**). To remove trail pheromones between assays, we covered the floor of the box with white poster board that we replaced between each assay. We first collected five foragers, defined as any worker outside the nest, from inside the colony container and placed them in a large petri dish. We placed the petri dish upside-down in the middle of a circular arena in the center of the filming box and waited five minutes to give the workers time to settle down after being handled. After the five minutes, we gently removed the petri dish so the workers were free to move around the arena and filmed the workers exploring the arena for 15 minutes.

Next, we replaced the poster board inside the filming box and placed the five foragers, all remaining foragers from inside the colony container, and the nest containing the rest of the workers, queens, and brood inside a petri dish. We placed the petri dish containing the entire colony upside-down in the center of the arena and waited five minutes before lifting the petri dish and filming for 15 minutes.

We analyzed the videos of the five foragers using custom made tracking software (https://github.com/swarm-lab/trackR; accessed 2017) to track the location of each ant in each frame of the video. To avoid the effect of the arena wall on ant trajectories, we removed all tracks where the ants were within 3 mm of the wall, resulting in many separate trajectories within each video for each ant. Next, for each sub-trajectory, we calculated the net squared displacement (NSD) by taking the square of the distance traveled by each ant between the starting location and each successive location along the rest of the trajectory. To calculate the diffusion coefficient, we took the slope of the plot of NSD over time and fit the equation:

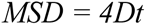

where mean squared displacement (*MSD*) is the slope of NSD over time, *D* is the diffusion coefficient, and *t* is time (Börger & Fryxell 2012). The diffusion coefficient served as a measure of how quickly the ants collectively explored a novel space.

In addition, for both the five forager and entire colony videos, we calculated the arena coverage and coverage redundancy over time. First, we computed the absolute difference between each frame of the recorded video and a background image of the experimental setup without ants in it. When a pixel had a large absolute difference, it meant an ant was present on that pixel in a given frame. We then applied a threshold to the difference image and classified all the pixels with a difference value above the threshold as “ant-covered” pixels and gave them a value of 1, and all the pixels with a difference value below the threshold as “background” pixels and gave them a value of 0. Finally, we computed the cumulative sum of the segmented images over time and calculated for each of them the arena coverage as the percentage of the pixels with a value of at least 1 (i.e. what fraction of pixels have been visited by ants at least once; **Figure 1)**.

**Figure 1.**
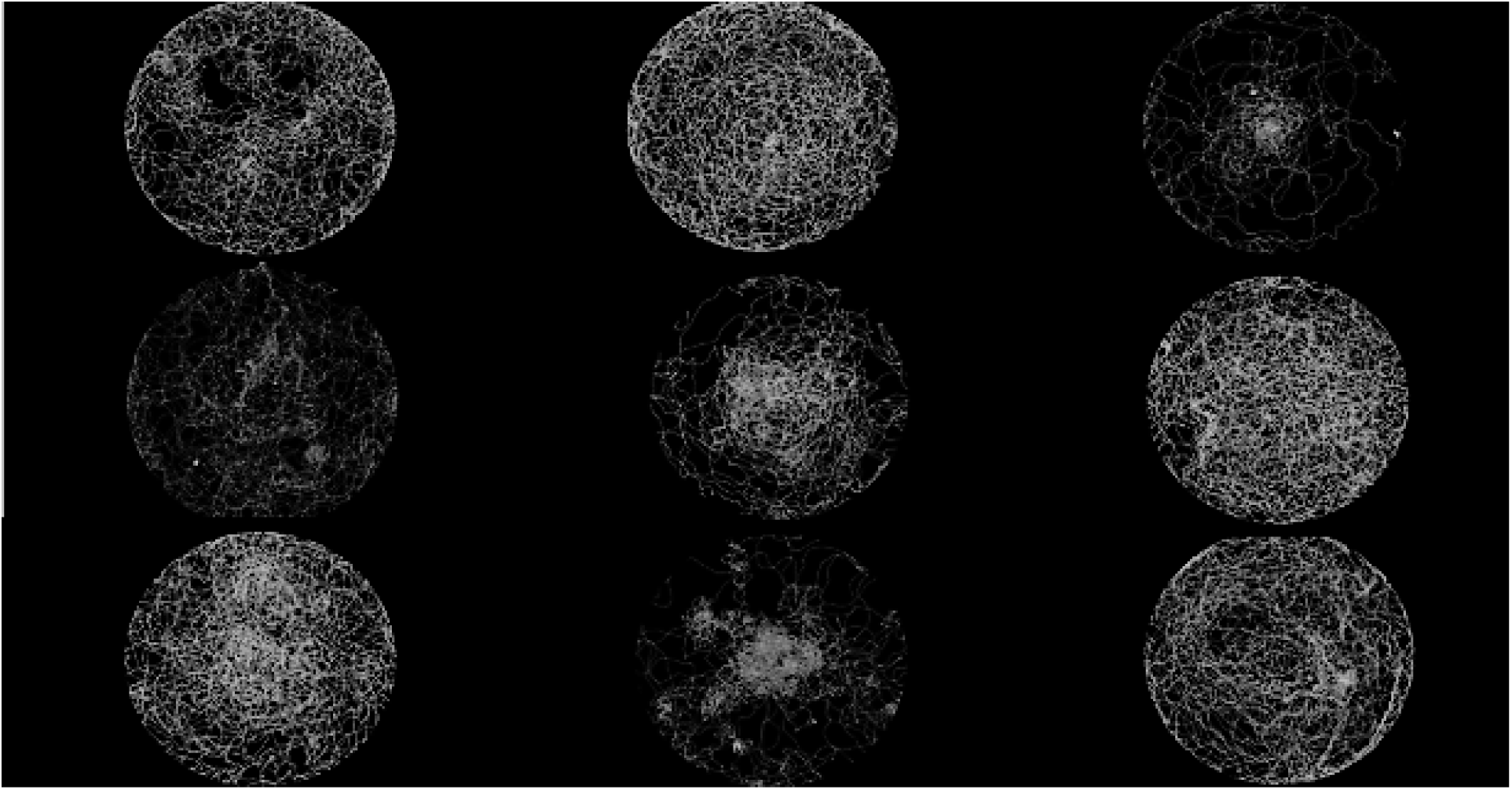
Nine representative plots showing variation among colony genotypes in the exploratory patterns of groups of five foragers. The plots show the tracks (white pixels) of the ants as they explore a novel arena.

We will refer to three exploratory behaviors as “exploratory rate”, “group exploration”, and “colony exploration”. “Exploratory rate” refers to the diffusion coefficient of groups of five ants, “group exploration” to the percent of the arena covered by the groups of five foragers, and “colony exploration” to the percent of the arena covered by the entire colony.

#### (ii) Foraging assay

We conducted the foraging assay on each colony replicate the day after the exploratory assay and after the colony replicates had been starved for a week. We melted the agar-based synthetic diet and soaked a cotton ball in the liquid. When the cotton ball solidified, we placed it on the plateau of a 3D printed ramp and placed the ramp inside a colony container on the opposite site of the nest (**Supplemental figure 2B**). Once an ant first discovered the food, we started filming and filmed for one hour. If no ant discovered the food in 30 minutes, we started the recording. We calculated the foraging rate by manually counting the number of ant visits to the plateau of the ramp in each video. Because many ants went back and forth from the food to the nest, we counted many ants more than once.

#### (iii) Aggression assay

Like other unicolonial ant species, *M*. *pharaonis* workers show little to no aggression towards *M*. *pharaonis* workers from other colonies (Schmidt et al. 2010). To get *M*. *pharaonis* workers to act aggressively, and to be able to quantify aggression against a constant “enemy” for all of our experimental colonies, we used workers from a single *Monomorium dichroum* colony that had been kept in the lab under the same conditions as the *M*. *pharaonis* colonies for 5 years. We conducted the aggression assays a week after the foraging assays. We first collected twenty foragers of both species and placed them in separate small petri dishes (**Supplemental figure 3**). We placed both small petri dishes upside down in a large petri dish for five minutes before lifting both petri dishes and allowing the workers of both species to interact. Every 5 minutes for one hour, we manually counted and recorded the number of *M*. *pharaonis* workers that were biting *M*. *dichroum* workers. We defined aggression as the average number of *M. pharaonis* workers biting *M. dichroum* workers across all observations within an hour. We froze all of the ants used in the aggression assay so that we did not reuse *M*. *dichroum* workers in more than one assay.

### (c) Colony productivity and body mass measurements

As a measure of colony productivity, we surveyed each colony replicate once per week and counted the number of workers and brood at all developmental stages. *M*. *pharonis* colonies usually only produce new gynes (virgin queens) and males in the absence of fertile queens (Edwards 1991; Warner et al. 2018). Therefore, in order to induce the production of new gynes and males, we removed queens at the start of the fifth week, after the aggression assay. We conducted weekly surveys until all brood matured into worker, gyne, or male pupae. In addition to colony productivity data for the total number of workers, gynes, and males produced, the weekly surveys also allowed us to calculate colony caste and sex ratio. We defined caste ratio as the number of gynes relative to the total number of females produced, and sex ratio as the number of gynes relative to the total number of reproductives (gynes and males) produced. To measure body size, we collected 15 worker pupae, 10 gyne pupae, and 10 male pupae from each colony replicate. We dried the pupae out in a drying oven for 24 hours before weighing.

### (d) Heritability and genetic correlation analysis

We performed all statistical analyses in R version 3.4.1 (R Core Team 2014). We estimated the repeatability of all measured phenotypes across colony replicates using a generalized linear mixed model (GLMM) approach in the R package MCMCglmm (Hadfield 2010). We included block as a random factor to account for the fact that the samples were collected at different time points from the replicate colonies and included colony identity as a random effect and *Wolbachia* infection status as a fixed effect (two of the original eight lineages included in the heterogeneous stock were infected with *Wolbachia;* Schmidt et al. 2010, Pontieri et al. 2017).

To estimate the heritability of, and genetic correlations between, all measured phenotypes, we used an animal model approach. Animal models estimate genetic parameters of traits by evaluating how patterns of observed phenotypic covariance between all pairs of individual “animals” is predicted by the expected genetic relatedness between individuals, based on pedigree (Kruuk 2004, de Villemereuil 2012). For our study, “individual animals” were replicate colonies, the pedigree was the known pedigree across nine generations of the *M. pharaonis* colonies in our mapping population, and the pedigree specifically represented genealogical relationships among the workers (i.e. the worker offspring of queen and male parents) that make up the replicate colonies of the mapping population. We thus assessed the degree to which the expected genotype of workers predicted the observed collective behavior or group-level phenotype measured for groups of workers from replicate colonies. Note that while we focused only on how expected worker genotype was associated with variation in worker collective behavior and colony-level traits, it is certainly possible that the genotypes of other types of colony members (i.e. queens or sibling larvae) also contributes to variation in the group-level traits we measured. Such effects can be independently estimated as described above, if very large datasets are available (e.g., Brascamp et al. 2016 used a honey bee dataset with 15,000 colonies), or alternatively, these effects can be experimentally teased apart with cross-fostering (Linksvayer 2006, 2007, Linksvayer et al. 2009). However, we did not have enough power in our dataset to separately estimate potential queen genetic effects, and effects of larval genotype are always completely confounded with worker genotype barring experimental cross-fostering.

Specifically, we used the R package MCMCglmm to run animal models using a Bayesian Markov chain Monte Carlo (MCMC) approach (de Villemereuil 2012). We accounted for the fact that ants are haplodiploid (males are haploid, females are diploid) by constructing the pedigree as if the traits were all sex-linked (Hedrick & Parker 1997). We used weakly informative priors for 1,000,000 iterations, with a burn-in period of 10,000 iterations and stored estimates every 500 iterations (full R script included in supplemental material; following de Villemereuil 2012). We assessed convergence of the models by visually inspecting estimate plots and assessing the autocorrelation values (de Villemereuil 2012). We analyzed whether behaviors were phenotypically correlated with each other (i.e. behavioral syndromes) using Spearman rank correlations and corrected for multiple comparisons by using the “FDR” method in the R function “p.adjust.”

In our initial heritability estimates, we ignored two complications in our pedigree. First, between our conducting new crosses to produce new generations, our colonies went through multiple rounds of intranidal mating: when the fecundity of current queens declines, *M. pharaonis* colonies produce new gynes and males which stay in the nest and mate with each other (Berndt & Eichler 1987). Second, when a colony was the mother/father colony to multiple offspring colonies, we initially treated those offspring colonies as half siblings. However, because *M. pharaonis* colonies contain multiple queens, the new gynes and males they produce may be better thought of as cousins. To test whether either of these complications would affect our heritability estimates, we constructed multiple pedigrees and re-ran the heritability analyses. We constructed pedigrees in which one or two generations contained two rounds of intranidal mating, one or two generations considered reproductives from the same colony as cousins, and two generations of both intranidal mating and considering reproductives from the same colony as cousins.

### (e) Selection analysis

We defined fitness in two ways, as either the production of new reproductives (gynes or males) or new workers, and ran separate models for each fitness definition. In nature, *M. pharaonis* colonies reproduce by budding (i.e. new colonies are not founded independently by queens; Buczkowski & Bennett 2009), but instead, a number of queens and workers disperse with brood to form a new colony. Both new reproductives and new workers determine the growth rate and potential to bud for existing colonies, and hence are appropriate measures of colony fitness. We estimated the strength of selection using a multivariate standardized selection gradient approach as described by Morrissey & Sakrejda (2013). This method is similar to the approach outlined by Lande and Arnold (1983) and uses spline-based generalized additive models to model the relationship between fitness and traits. We normalized all behaviors to a mean of zero and a standard deviation of one so that the selection estimates represent standardized values (Lande & Arnold 2983; Morrissey & Sakrejda 2013). We included all five behaviors and block in all models and estimated selection gradients and prediction intervals after 1000 bootstrap replicates (Morrissey & Sakrejda 2013).

## Results

### (a) Repeatability and heritability estimates

All five behaviors, caste and sex ratio, and worker, gyne, and male body mass were significantly repeatable across replicate colonies (**Supplemental table 1**). We estimated the heritability of the five collective behaviors to be between 0.17 and 0.32, with a median value of 0.21 (**Figure 2)**. We estimated the heritability of worker body mass to be 0.34, gyne body mass to be 0.46, and male body mass to be 0.53 (**Figure 2)**. We estimated the heritability of five colony productivity measures to be between 0.001 and 0.46, with a median value of 0.24 (**Figure 2**). Finally, we estimated the heritabilty of colony caste and sex ratio to be 0.26 and 0.23, respectively (**Figure 2)**.

**Figure 2.**
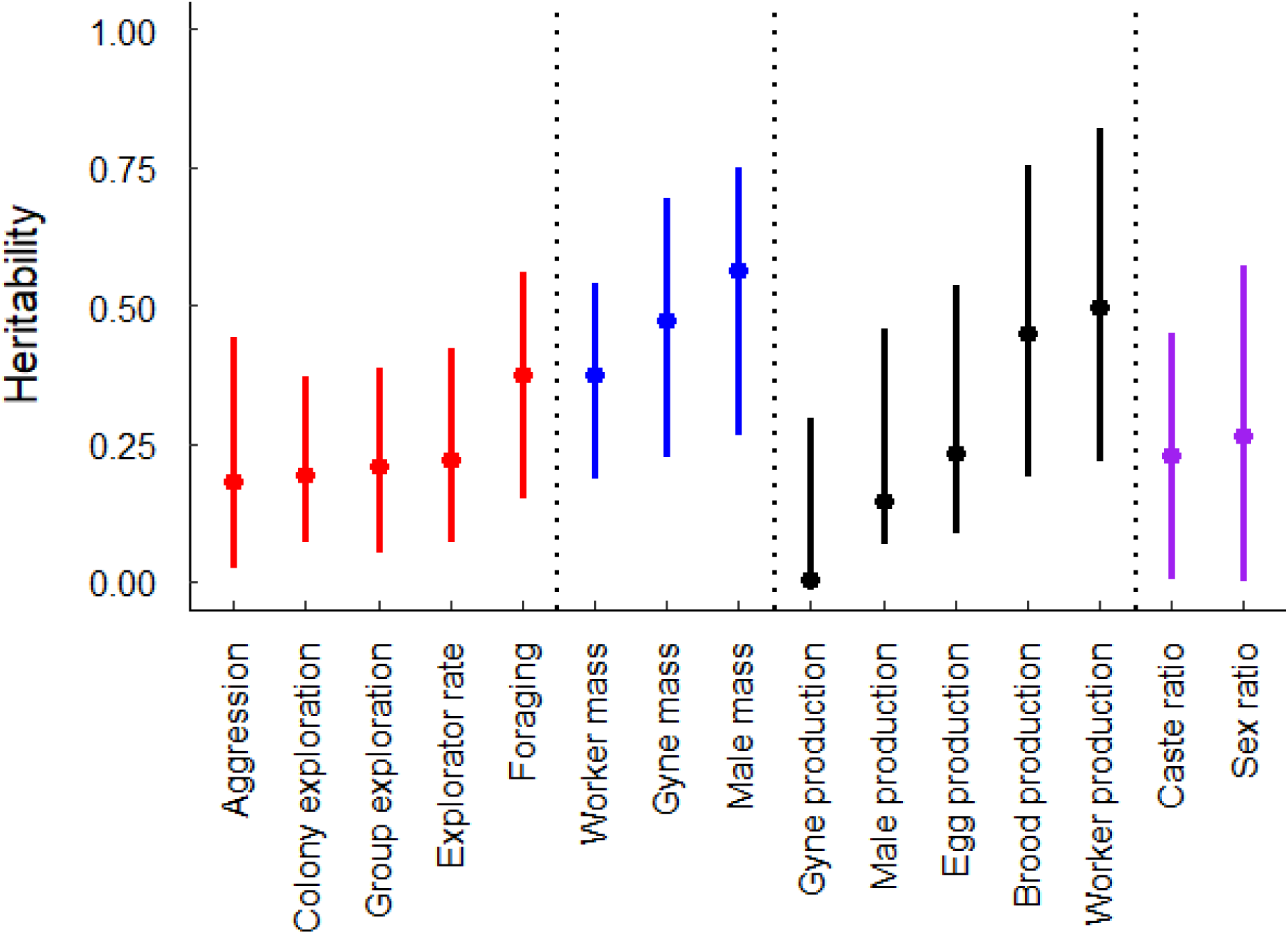
Caterpillar plot of heritability estimates +/− 95% confidence intervals grouped by category. Collective behaviors (red), body mass (blue), colony productivity (black), and caste and sex ratio (purple) are designated by different colors.

We compared our initial heritability estimates with heritability estimates using five different modified pedigrees that considered intranidal mating and/or considering offspring colonies as cousins rather than half siblings. The difference between the initial heritability estimates and the estimates when using the five modified pedigrees were small, less than 0.1 for all phenotypes except sex ratio, which differed by up to 0.24 (**Supplemental table 2**).

### (b) Phenotypic and genetic correlation estimates

We found phenotypic correlations among the five measured collective behaviors (**Figure 3**). Foraging rate was negatively correlated with aggression and positively correlated with both group exploration and colony exploration. Aggression was negatively correlated with exploratory rate. Group exploration and colony exploration were positively correlated. The genetic correlation estimates ranged from −0.05 to 0.17 but the 95% CIs all overlapped with zero (see **Figure 3** and **Supplemental table 3** for estimates and 95% CI). The genetic correlation estimates between behaviors and all other traits, as well as among all the other traits, were mostly small and all had 95% CI that overlapped with zero (**Supplemental table 4**).

**Figure 3.**
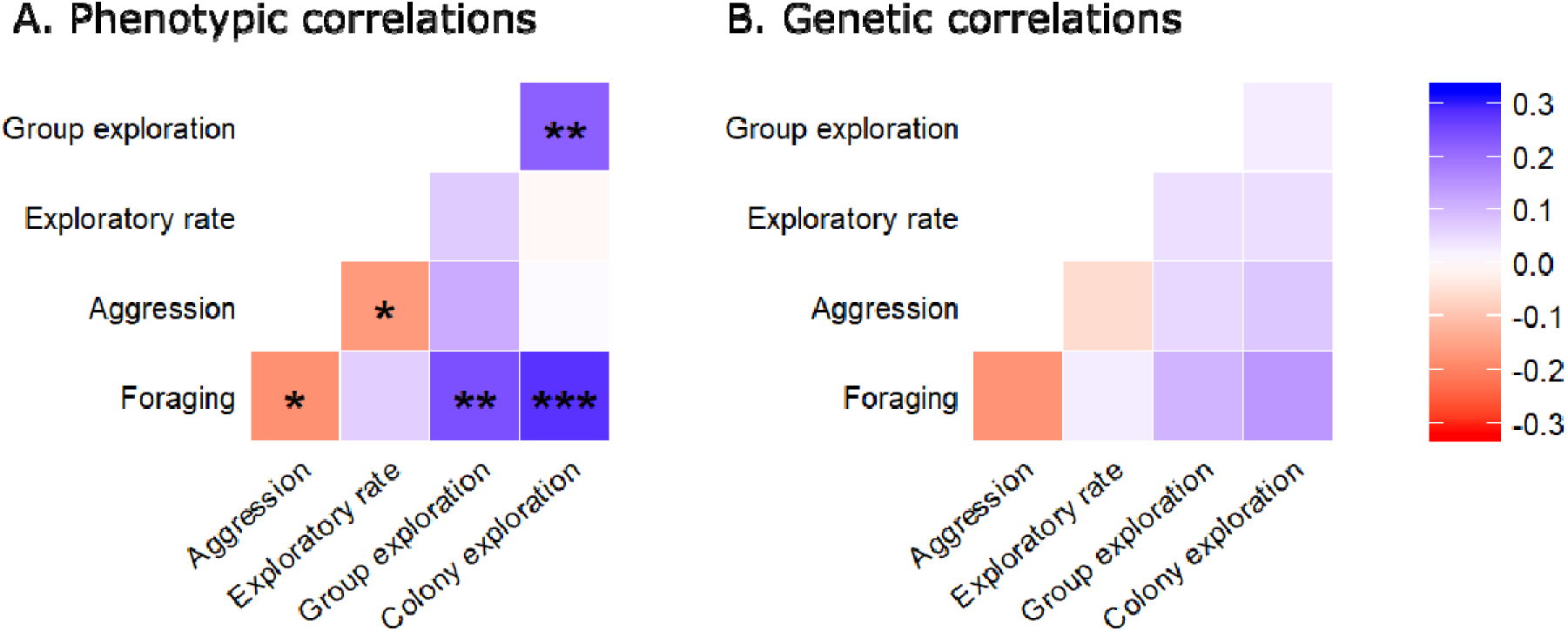
Heatmaps showing phenotypic (A) and genetic (B) correlations between collective behaviors. For the phenotypic correlations, asterisks within cells correspond to p values (adjusted for multiple comparisons; p < 0.05 = *; p < 0.01 = **; p < 0.001 = ***; no symbol indicates p > 0.05) and the colors correspond to the magnitude and sign of the Spearman rank correlation coefficient. None of the genetic correlations were significant (all 95% CIs overlapped with zero).

### (c) Selection gradients

When defining fitness as the number of reproductives (gynes + males) produced by the colony, we found evidence for positive linear selection on foraging and negative linear selection on exploratory rate (**Table 1, Figure 4**). We found no evidence for quadratic selection. When defining fitness as the number of workers produced by a colony, we found evidence for positive linear selection on foraging and no evidence for quadratic selection (**Table 1, Figure 4**). When defining fitness as either the production of new reproductives or workers, we found no evidence for correlational selection between any of the five behaviors. Finally, we found no evidence for linear or quadratic selection on worker, gyne, or male body mass (**Supplemental table 5**).

**Figure 4.**
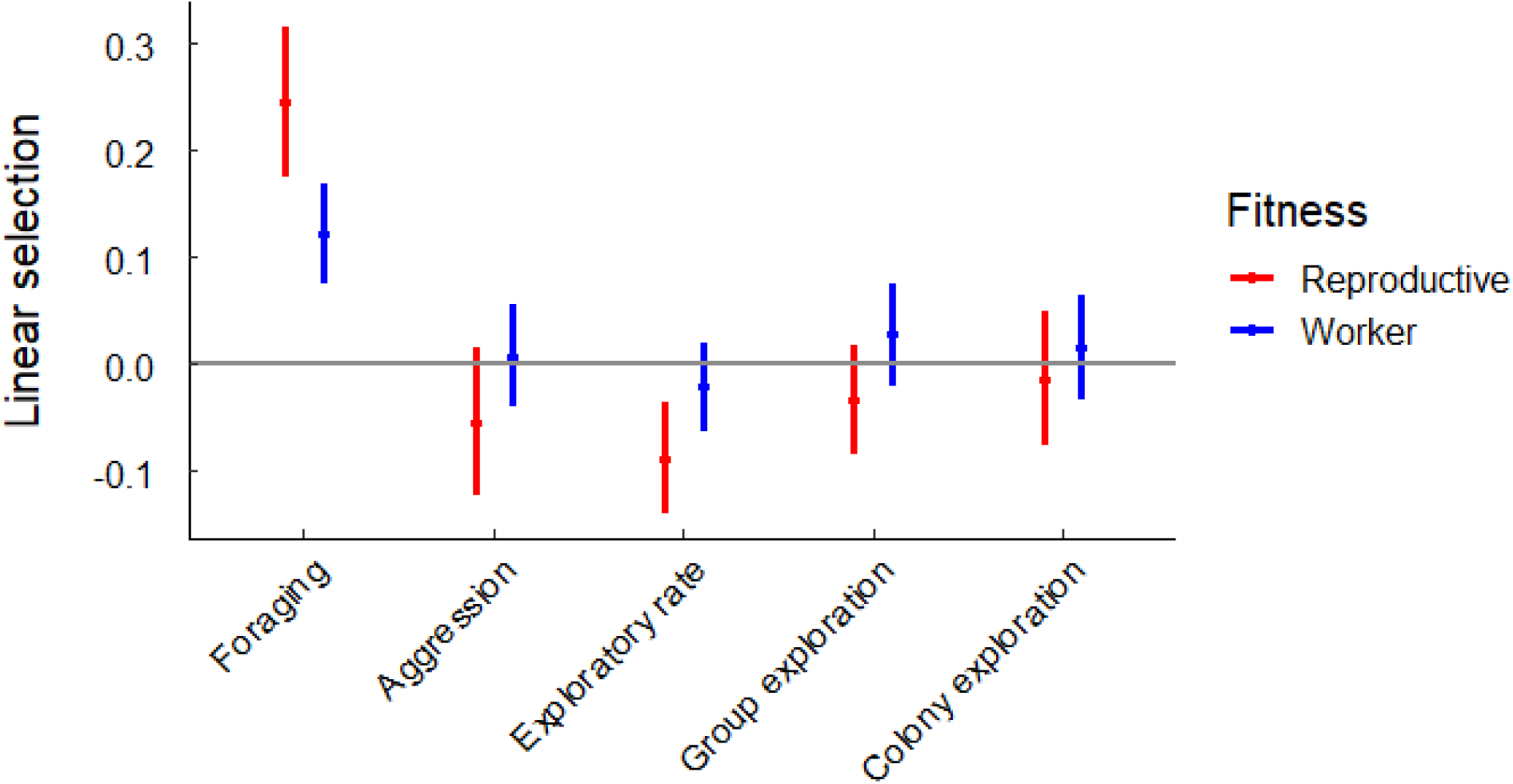
Caterpillar plot showing linear (a) and quadratic (b) selection gradients +/− SE. Asterisks indicate estimates that are significant. Positive values indicate directional selection for increased trait values (e.g., there is directional selection for increased foraging rate when fitness is measured by the number of reproductives or workers produced) and negative values indicate directional selection for decreased trait values (e.g., there is directional selection for decreased exploratory rate when fitness is measured as the number of reproductives produced).

**Table 1.**
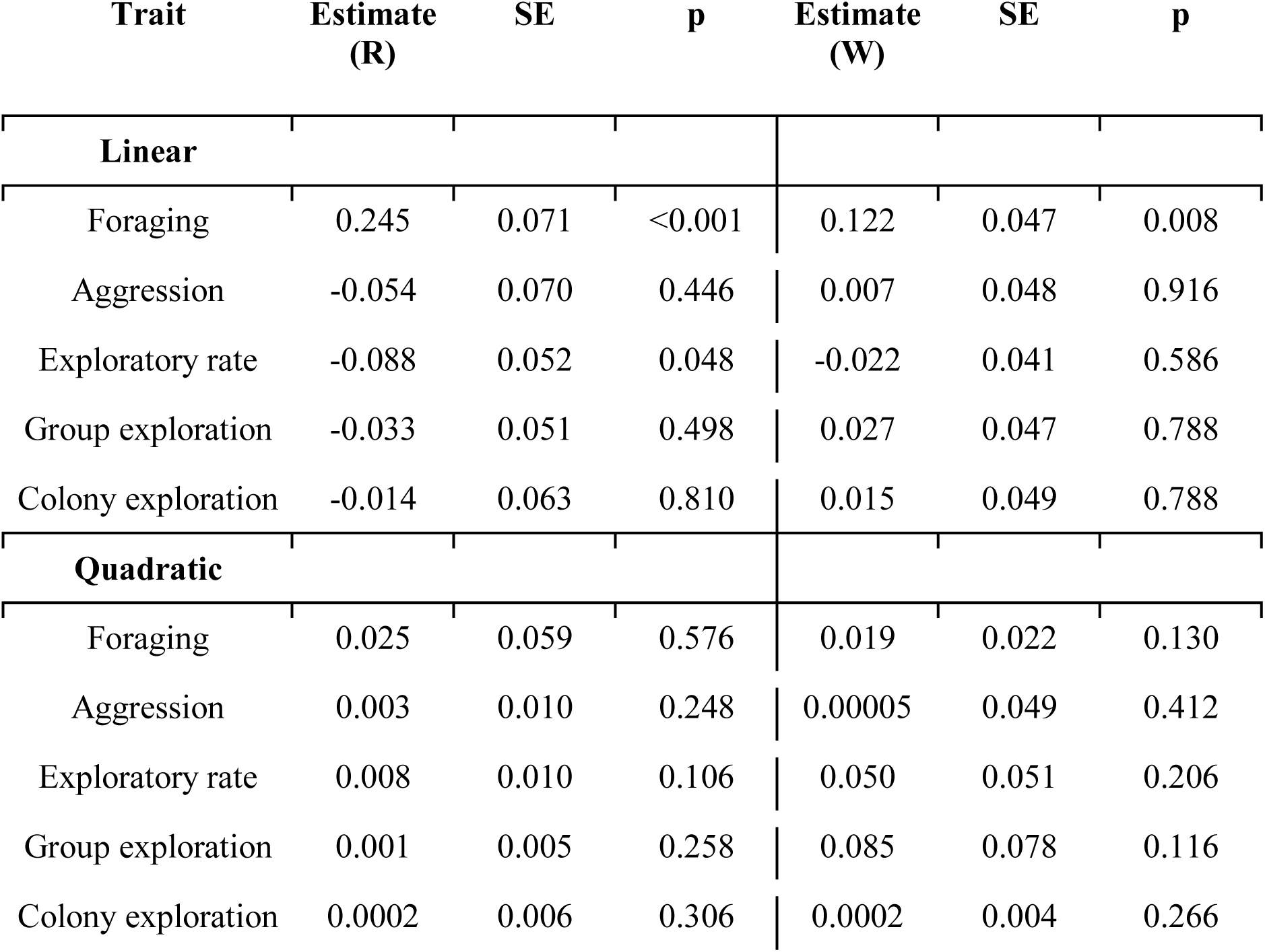
Linear and quadratic selection estimates for behaviors using either reproductive (R) or worker (W) production as the measurement of fitness.

| Trait | Estimate<br>(R) | SE | p | Estimate<br>(W) | SE | p |
| --- | --- | --- | --- | --- | --- | --- |
| **Linear** |  |  |  |  |  |  |
| Foraging | 0.245 | 0.071 | <0.001 | 0.122 | 0.047 | 0.008 |
| Aggression | -0.054 | 0.070 | 0.446 | 0.007 | 0.048 | 0.916 |
| Exploratory rate | -0.088 | 0.052 | 0.048 | -0.022 | 0.041 | 0.586 |
| Group exploration | -0.033 | 0.051 | 0.498 | 0.027 | 0.047 | 0.788 |
| Colony exploration | -0.014 | 0.063 | 0.810 | 0.015 | 0.049 | 0.788 |
| **Quadratic** |  |  |  |  |  |  |
| Foraging | 0.025 | 0.059 | 0.576 | 0.019 | 0.022 | 0.130 |
| Aggression | 0.003 | 0.010 | 0.248 | 0.00005 | 0.049 | 0.412 |
| Exploratory rate | 0.008 | 0.010 | 0.106 | 0.050 | 0.051 | 0.206 |
| Group exploration | 0.001 | 0.005 | 0.258 | 0.085 | 0.078 | 0.116 |
| Colony exploration | 0.0002 | 0.006 | 0.306 | 0.0002 | 0.004 | 0.266 |

To further put our results into context, we estimated the proportion of variance among our colonies for both measures of fitness (the productions of new reproductives and workers) that was explained by variation in any of our five behavioral variables, experimental block, or *Wolbachia* infection status. For the production of new reproductives, we found that aggression explained the largest amount of the variance (5.29%), followed by foraging (2.29%), group exploration (1.94%), exploratory rate (0.52%), and colony exploration (0.33%) (**Supplemental table 6**). For the production of new workers, we found that foraging explained the largest amount of the variance (1.29%), followed by aggression (0.53%), colony exploration (0.34%), group exploration (0.27%), and exploratory rate (0.08%) (**Supplemental table 6**).

## Discussion

Collective behavior is ubiquitous in nature and presumed to have strong fitness consequences for group members. Moreover, repeatable variation in collective behavior (often described as collective or group-level “personality”) has been commonly observed (Bengston & Jandt 2014; Planas-Sitjà et al. 2015; Jolles et al. 2017; Wright et al. 2019). However, little is known about the heritability or genetic architecture of collective behavior and how collective behavior is shaped by selection. A major difficulty for elucidating the genetic basis of collective behavior is that, unlike individual-level behavior, collective behavior by definition depends on social interactions among members of the group. As a result, the genetic architecture of collective behavior fundamentally depends on the collective genetic make-up of these individuals (McGlothlin et al. 2010; Linksvayer 2006, 2015). Quantifying patterns of selection on group-level traits also has an added level of difficulty because the level of replication is the group (e.g., colony) and not the individual. Here, we begin to elucidate the genetic architecture underlying collective behavior and other group-level traits and to characterize how selection acts on these traits in a laboratory population of the ant *Monomorium pharaonis* that we created for this purpose. We provide evidence that variation in collective behaviors, including foraging, aggression, exploratory rate, group and colony exploration, and other group-level traits measured in the laboratory is heritable, phenotypically and genetically correlated, and shaped by selection.

We estimated the heritability of collective behaviors to be between 0.22 and 0.40, which was generally lower than the heritability estimates for body size (0.38 to 0.58), colony productivity (0.14 to 0.75), and caste (0.42) and sex ratio (0.49) (**Figure 2**, also see **Supplemental table 2** for heritability estimates using more complex pedigrees). These heritability estimates demonstrate that all of the phenotypes we measured, including collective behaviors, have the ability to respond to short term selection on standing genetic variation. Although numerous studies have examined the genetic architecture of group-level traits in honey bees (Rinderer et al. 1983; Collins et al. 1984; Milne 1985; Moritz et al. 1987; Bienefeld & Pirchner 1990; Pirchner & Bienefeld 1991; Harris & Harbo 1999; Boecking et al. 2000; Hunt et al. 2007), this is one of the first studies to examine the genetic architecture or the evolution of collective behavior and other group-level traits in an ant species.

Although our heritability estimates are somewhat higher than other estimates of heritability across animal taxa (e.g. the heritability of individual-level behaviors was on average 0.14; Dochtermann et al. 2015), heritability estimates can vary widely, and all-else-equal are expected to be higher in animals bred in captivity than in nature because environmental conditions in the laboratory are controlled (Simmons & Roff 1994). Furthermore, the heritability estimates for all of our measured group-level phenotypes may be higher than individual-level behaviors because the heritability of traits influenced by social interactions includes the contribution of heritable components of the social environment (Linksvayer 2006; Bijma et al. 2007a, 2007b; Linksvayer et al. 2009; McGlothlin et al. 2010; Bijma 2011). There is ample empirical and theoretical evidence that this form of “hidden heritability” contributes to the heritable variation and also the evolutionary response to selection for social traits (Wade 1976; Moore 1990; Muir 2005; Linksvayer 2006; Bijma et al. 2007b; Bergsma et al. 2008; Wade et al. 2010; Bijma 2011). Because we kept all components of the social environment intact across replicate sub-colonies of each colony genotype (i.e. the workers, queens and brood were all from the same parent colony), our heritability estimates do not partition out the relative contributions of variation in the workers’ own genomes from variation in the genomes of other colony members (Linksvayer 2006; Linksvayer et al. 2009).

We found evidence for both phenotypic and genetic correlations between collective behaviors. Suites of phenotypically correlated behaviors are termed “behavioral syndromes” and have been documented throughout the animal kingdom, including in social insects (Sih et al. 2004; Jandt et al. 2014). The behavioral syndrome we found in *M. pharaonis* consisted of a positive correlation between foraging and exploration, which were both negatively correlated with aggression. Our phenotypic and genetic correlation estimates were generally similar. For example, the four strongest genetic correlation estimates (Foraging - Aggression; Foraging - Forager coverage; Foraging - Colony coverage; Forager coverage - Colony coverage; **Figure 3**) were also four of the five significant phenotypic correlations and were all in the same direction. However, our genetic correlation estimates were generally very weak (i.e. not significantly different than zero) and only one of our genetic correlation estimates was bound away from zero (the correlation between foraging and colony exploration).

Traditionally, behavioral ecologists relied on the assumptions that all behavioral traits were heritable, not strongly genetically correlated, and thus free to evolve independently from other traits in response to patterns of selection on each trait. This approach was termed the “phenotypic gambit” (Grafen 1984). Our results generally support these assumptions as we found moderate estimates of heritability for all five behavioral variables and relatively weak genetic correlation estimates. These results suggest that collective behaviors are free to respond to selection, and that the underlying genetic architecture will not constrain long-term optimization by natural selection (Lynch & Walsh 1998; Wilson et al. 2010).

We calculated the strength and direction of selection acting on collective behavior and found evidence for both positive and negative linear selection (**Figure 4, Table 1**). The absolute value of our estimates of the strength of linear and quadratic selection are similar or slightly smaller than estimates of selection in wild populations (Kingsolver et al. 2001). The strongest pattern of linear selection we found was for foraging, indicating that colonies with higher foraging rates produced more reproductives as well as workers. A higher foraging rate is presumably associated with higher input of resources for the colony, allowing colonies to be more productive.

We conducted the current study in a laboratory environment, which enabled us to strictly control the demographic make-up (i.e. queen number, worker number, etc.) and precise environmental conditions experienced by the three colony replicates for each of our 81 colony genotypes. Such control in particular is valuable given the complexity of social insect colonies (Linksvayer 2006; Kronauer & Libbrecht 2018). However, we also acknowledge the caveat that our choice to conduct our study in a controlled laboratory environment likely had strong effects on both our estimates of heritability and genetic correlations, as well as our estimates of the pattern and magnitude of selection. In particular, it is difficult to know how the fitness consequences of variation in collective behavior that we observed would change in a more natural setting. One possibility is that we might observe positive linear selection for aggression since aggression has no obvious benefit in the lab but may have benefits in nature. Laboratory-based estimates of natural selection are commonly used to test predictions of evolutionary theory (Fuller et al. 2005). For example, researchers used lab-based manipulations to test predictions of density-dependent selection theory in *Drosophila melanogaster* (e.g. Mueller 1997; Dasgupta et al. 2019). Our study provides some of the first evidence that natural selection can shape collective phenotypes, an assumption that is rarely tested, on a scale that is likely not feasible in a field study. Furthermore, because *M*. *pharaonis* tends to be found in association with humans, both in the tropics in their presumed native range (Wetterer 2010) and in heated buildings in introduced temperate regions, the laboratory conditions of our study might be more similar to the “natural” conditions experienced by our study species than other non-synanthropic species.

Overall, this study increases our understanding of the genetic architecture of collective behavior and demonstrates that it is strongly shaped by natural selection. Future studies should focus on identifying the mechanisms by which genes function to influence collective behavior and how variation in these genes affects patterns of variation for collective behavior within populations. Candidate gene approaches have been used successfully to demonstrate the roles of the ant ortholog of the *foraging* gene (Ingram et al. 2005; Lucas & Sokolowski 2009; Ingram et al. 2016; Bockoven et al. 2017; Page et al. 2018) and dopamine (Friedman et al. 2018). In addition to candidate gene approaches, future studies should utilize unbiased approaches such as quantitative trait locus (QTL) mapping in mapping populations (e.g., Hunt et al. 1998; 2007) such as ours, and association mapping in natural populations (e.g., Kocher et al. 2018). Additionally, future research should aim to understand the mechanisms underlying the expression of collective behavior (Friedman et al. 2019). For example, chemical communication (e.g. cuticular hydrocarbons, pheromones) likely plays a large role in regulating collective behavior in social insects. Finally, future studies should seek to disentangle the contribution of workers’ own genomes and the composite sociogenome of their nestmates (including other workers, queens, and brood), by using cross-fostering approaches and experimentally setting up mixed worker groups (Morowitz & Southwick 1987; Calderone & Page 1992; Linksvayer 2006; Linksvayer et al. 2009; Gempe et al. 2012).

