## Supplemental files for "Ant collective behavior is heritable and shaped by selection"

**Supplemental material**

**Supplemental table 1.** Repeatability estimates across replicates

|  | **R** | **Low 95 CI** | **High 95 CI** |
| --- | --- | --- | --- |
| Foraging | 0.460 | 0.307 | 0.569 |
| Aggression | 0.530 | 0.412 | 0.655 |
| Exploratory rate | 0.505 | 0.364 | 0.635 |
| Group exploration | 0.297 | 0.144 | 0.414 |
| Colony exploration | 0.433 | 0.321 | 0.603 |
| Caste ratio | 0.475 | 0.305 | 0.588 |
| Sex ratio | 0.674 | 0.553 | 0.748 |
| Worker mass | 0.510 | 0.438 | 0.678 |
| Gyne mass | 0.687 | 0.557 | 0.763 |
| Male mass | 0.577 | 0.326 | 0.711 |

**Supplemental table 2.** Heritability estimates for all traits using six different pedigrees.

**Supplemental table 3.** Genetic correlation estimates among behaviors +/- 95% confidence intervals.

|  | **Correlation** | **Low 95 CI** | **High 95 CI** |
| --- | --- | --- | --- |
| Foraging - Aggression | -0.166 | -0.458 | 0.133 |
| Foraging - Exploratory rate | 0.024 | -0.218 | 0.223 |
| Foraging - Group exploration | 0.097 | -0.250 | 0.285 |
| Foraging - Colony exploration | 0.135 | -0.140 | 0.324 |
| Aggression - Exploratory rate | -0.055 | -0.262 | 0.173 |
| Aggression - Group exploration | 0.051 | -0.257 | 0.318 |
| Aggression - Colony exploration | 0.072 | -0.192 | 0.254 |
| Exploratory rate - Group exploration | 0.041 | -0.188 | 0.220 |
| Exploratory rate - Colony exploration | 0.041 | -0.171 | 0.206 |
| Group exploration- Colony exploration | 0.023 | -0.143 | 0.245 |

**Supplemental table 4.** Genetic correlation estimates between behaviors and other traits +/- 95% confidence intervals.

**Supplemental table 5. Linear and quadratic selection estimates for worker, gyne, and male body mass using either reproductive (R) or worker (W) production as the measurement of fitness.**

| **Trait** | **Estimate**  **(R)** | **SE** | **p** | **Estimate**  **(W)** | **SE** | **p** |
| --- | --- | --- | --- | --- | --- | --- |
| **Linear** |  |  |  |  |  |  |
| Worker mass | -0.014 | 0.070 | 0.974 | 0.006 | 0.044 | 0.896 |
| Gyne mass | 0.029 | 0.080 | 0.980 | -0.005 | 0.050 | 0.938 |
| Male mass | 0.091 | 0.081 | 0.268 | -0.017 | 0.057 | 0.720 |
| **Quadratic** |  |  |  |  |  |  |
| Worker mass | <0.001 | 0.054 | 0.394 | <0.001 | 0.086 | 0.448 |
| Gyne mass | 0.013 | 0.070 | 0.364 | 0.064 | 0.063 | 0.264 |
| Male mass | 0.008 | 0.018 | 0.264 | <0.001 | 0.006 | 0.548 |

**Supplemental table 6. Proportion of the variance among colonies in the production of reproductives and workers explained by behavioral traits, experimental block, and *Wolbachia* infection status.**

| **Trait** | **Variance explained (%)**  **(Reproductives)** | **Variance explained (%)**  **(Workers)** |
| --- | --- | --- |
| Foraging | 2.29 | 1.29 |
| Aggression | 5.29 | 0.53 |
| Exploratory rate | 0.52 | 0.08 |
| Group exploration | 1.94 | 0.27 |
| Colony exploration | 0.33 | 0.34 |
| Block | 44.76 | 28.04 |
| *Wolbachia* | 1.19 | 1.00 |

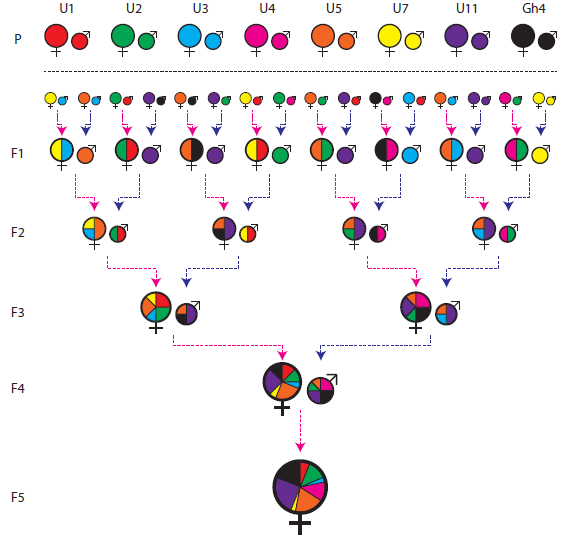

**Supplementary figure 1.** Crossing scheme employed for the creation of the *M. pharaonis* mapping population. For simplicity, only the crosses needed to create a generation F5 colony are shown, although eight parental colonies (P) were sequentially intercrossed for nine generations. The black dashed horizontal line separates generations of inbreeding and the onset of the crossing procedure. Briefly, gynes (♀) and males (♂) from each of the eight parental colonies (colour coded) were collected and crossed with sexuals from other parental colonies. From the resulting F1 offspring, new gynes and males were collected (pink and blue dashed arrows, respectively) and crossed in order to give birth to the F2 generation. The same protocol was applied for the subsequent generations. The expected genetic contribution of each parental colony to the genotype of each individual represented in the figure is indicated by different sizes of pie chart slices, colour coded according to parental colony.

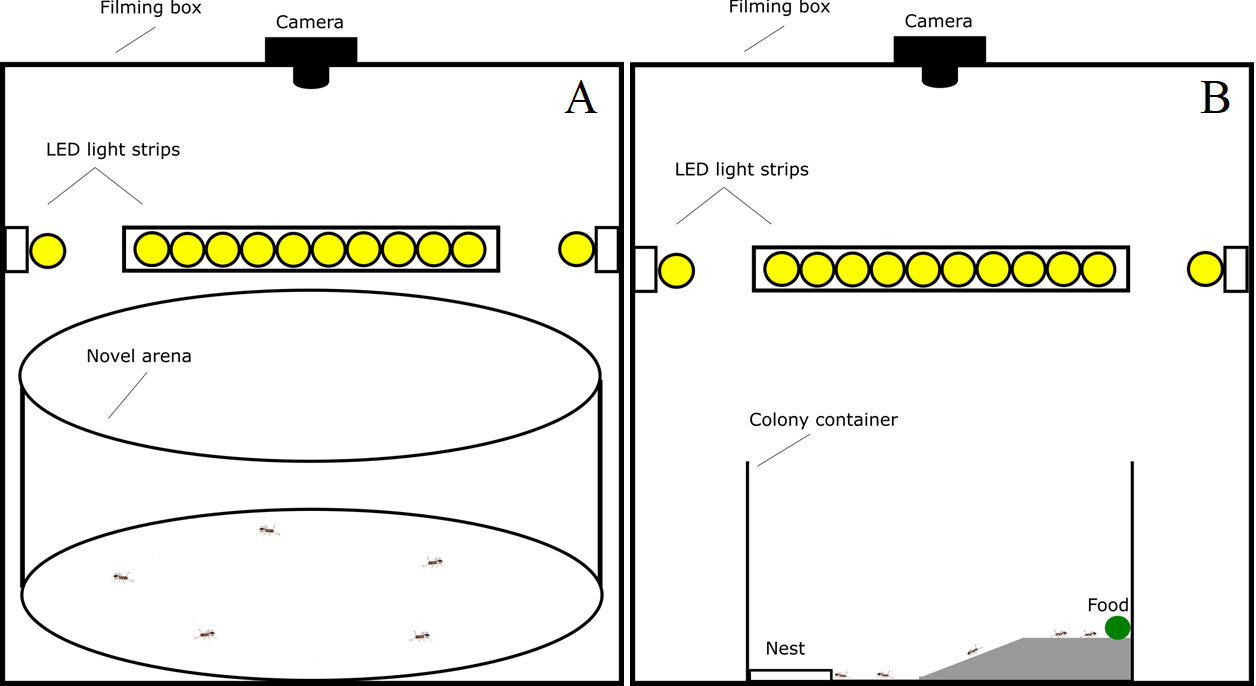

**Supplemental figure 2.** A diagram showing the setup of the exploratory and foraging assays. We conducted the assays inside a filming box with LED lights arranged along the walls and a camera mounted on the top to film the arena from above. In the exploratory assay (A), we covered the floor of the box with poster board that we replaced between each assay to remove trail pheromones. We first collected five foragers, defined as any worker outside the nest, from inside the colony container and placed them in a large petri dish (14 cm in diameter, 2 cm high). We placed the petri dish upside-down in the middle of a circular arena (28 cm in diameter, 10 cm high) in the center of the filming box. We waited five minutes to give the workers time to settle down after being handled. After the five minutes, we gently removed the petri dish so the workers were free to move around the arena and filmed the workers exploring the arena for 15 minutes. Next, we replaced the poster board inside the filming box and placed the five foragers, all remaining foragers from inside the colony container, and the nest containing the rest of the workers, the queens, and the brood inside a petri dish. We placed the petri dish containing the entire colony upside-down in the center of the arena and waited five minutes before lifting the petri dish and filming for 15 minutes. In the foraging assay (B), we placed a food-soaked cotton ball on the plateau of a 3D printed ramp (4.5 cm wide, 2 cm high; 10 cm in length: incline = 5 cm, plateau = 5 cm). We placed a colony container (8.5 cm x 10.5 cm x 10.5 cm) inside the filming box and put the ramp against the wall on the opposite side of the nest. All four sides of the filming box were lined with LED light strips. Once an ant first discovered the food, we started filming and filmed for one hour. If no ant discovered the food in 30 minutes, we started the recording.

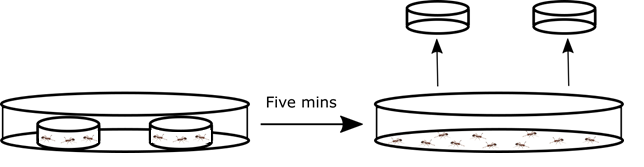

**Supplemental figure 3**. A diagram showing the setup of the aggression assay. We first collected twenty foragers of each species and placed them in separate small petri dishes (3.5 cm in diameter, 0.75 cm high). We placed both small petri dishes upside down in a large petri dish for five minutes before lifting both petri dishes and allowing the workers of both species to interact. Every 5 minutes for one hour, we recorded the number of *M*. *pharaonis* workers that were biting *M*. *dichroum* workers. We also recorded the number of workers of each species that were killed during the assay. Note: only 10 ants are shown in the diagram but 40 total ants (20 from each species) were used in each assay.
